# May a Strain Chlamydia Isolated From SARS Patient’s Autopsy Issues Inhibit the Proliferation of SARS-CoV? An Early Observation in Vitro

**DOI:** 10.1101/2021.08.27.457969

**Authors:** Xing Quan, Weifei Liu

## Abstract

We found that a strain chlamydia isolated from SARS patient’s autopsy issues could decrease the proliferation of SARS-CoV in vitro; The inhibitory factors distribute both in the extracellular and intracellular cultures.

## 0 Introduction

Interspecific competition is common in a ecosystem^[1, 2]^. The closer their ecological niche is, the more intense the competition is^[3, 4]^. Some species compete for resources through inhibiting the growth and reproduction of other species by releasing secondary metabolites or synthetic chemicals. For example, sage kills surrounding species by releasing volatile chemicals for competing surrounding nutritional resources, as well as walnuts (release juglone), we call it allelopathy^[5, 6]^; Another example is that penicillium secretes penicillin to inhibit the proliferation of Gram-positive bacteria^[7, 8]^. In our daily life, 70% - 80% of natural antibiotics are produced by actinomycetes^[9-13]^. Therefore, in the ecosystem, interspecific competition is very common to kill other competing species by releasing some chemicals.

We note that scrub-typhus can significantly inhibit the proliferation of human immunodeficiency virus type 1 (HIV-1, CXCR4) in HIV-1 patients. The HIV-1 viral load of two patients co-infected with Orientia Tsutsugamushi decreased below to the detection threshold^[14]^. Prof. OTOOLE,C found that the viral load of HIV-1 patients with mycoplasma infection also shown a decrease^[15]^; Other cases included that mycoplasma contamination inhibited the proliferation of Adenovirus and Mengovirus in their cell cultures^[16]^. These studies show that both rickettsia and mycoplasma can inhibit the proliferation of related different viruses under certain conditions. We note that rickettsia, mycoplasma and chlamydia all belong to the relationship between virus and bacteria, and because the ecological niche of chlamydia is closer to viruses compared to rickettsia and mycoplasma, it is doubtful whether chlamydia can inhibit the proliferation of viruses under certain conditions. A study conducted by Prof. Hong T et al showed that no Severe Acute Respiratory Syndrome Coronavirus (SARS-CoV) or few SARS-CoV were found in the tissue samples of some Severe Acute Respiratory Syndrome (SARS) patients, but a large number of chlamydia particles were found in their tissue samples^[17]^. Therefore, in order to investigate the relationship between SARS-CoV and the related strain chlamydia, We carried out the following series of experiments. This article is divided into four sections, the first section is materials and methods, the second section is results, the third section is discussions, and the final section is conclusions.

## 1 Materials and methods

### 1.1 Virus, chlamydia and cells

SARS-CoV (BJ01 strain), the strain Chlamydia-1 (isolated from the autopsy specimens of SARS-CoV patients), and Human Embryo Kidney 293 Cells were kindly provided by Prof. Hong T (Chinese Center for Disease Control and Prevention, Beijing, China); the strain *Chlamydia psittaci* (*C. psittaci*) was purchased from the China veterinary Culture Collection Center (No.CVCC2410). Virus and chlamydia stocks were prepared after virus inoculation into 293 cells respectively. Following incubation at 37°C for 48 h, cultured fluids were collected, clarified by low speed centrifugation and titrated for virus and chlamydia contents respectively.

For virus titration, serial dilutions of virus suspension were inoculated into cell culture plates and their fifty-percent Tissue Culture Infectious Dose (TCID_50_) titers determined using the method of Reed-Muench (1938). Finally, the viral titers were determined to 10^5.67^TCID_50_/ml. For chlamydia titration, 50 uL chlamydia suspension were inoculated in 293 cells with 1×10^5^ cells each well (a 13 mm round coverslip was paved in each well in advance). After incubation at 37°C for 48 h, the cells were fixed in absolute methanol for 10 min, and then stained by Giemsa method for 45 min, the number of inclusion-forming units (IFUs) of chlamydia were calculated through inverted microscope (Olympus, Japan). The final titration of the strain Chlamydia-1 were 1.2×10^5^ IFUs/mL, and the final titration of the strain *C. psittaci* were 3.0×10^5^ IFUs/mL.

### 1.2 Co-infection of SARS-CoV and chlamydia in 293 cells

For the co-infection of SARS-CoV and the strain Chlamydia-1: 293 cells were cultured in 200 uL dulbecco’s modified eagle’s medium (DMKM) supplemented with 10% fetal bovine serum (FBS) and 1% antibiotics/antimycotic, seeded in a 96-well cell plate with 1×10^5^ cells per well (a 13 mm round coverslip was paved in each well in advance), and then kept in 5% CO_2_, saturated humidity, and 37°C. While cell density grown to about 80%, it began to inoculate virus and chlamydia. Infection groups were divided into six groups, including SARS-CoV single infection group (A_1_ group), the strain Chlamydia-1 single infection group (B_1_ group), SARS-CoV and the strain Chlamydia-1 simultaneous infection group (C_1_ group), SARS-CoV infection first, then the strain Chlamydia-1 infection group (D_1_ group, the latter inoculation in 2 days later), the strain Chlamydia-1infection first, then SARS-CoV infection group (E_1_ group, as above) and control group (F_1_ group). when each inoculation, gently shake the cell plate every 20min for 1 hour, and then discard inoculated fluids, wash with 10% phosphate buffered saline (PBS). Finally, add 200 uL 10% FBS DMKM to continue to incubate those inoculated cells. Both the inoculation volume of virus and chlamydia were 50 uL. The viral load of SARS-CoV [by quantitative real-time polymerase chain reaction (qRT-PCR)] and the number of IFUs (by Giemsa method and inverted microscope) would be detected on the 3rd, 4th, 5th, 6th, 7th, 8th and 9th day after the first inoculation. The concentration of SARS-CoV would be diluted to 10^3.67^ TCID_50_/mL for the inoculation and the concentration of the chlamydia-1 strain were diluted 6.0×10^4^ IFUs/mL for the infection.

For the co-infection of SARS-CoV and the strain *C. psittaci* : The same as above, except replacing the strain Chlamydia-1 as the strain *C. psittaci*. The corresponding groups were numbered as A_2_, B_2_, C_2_, D_2_, E_2_, F_2_ respectively.

### 1.3 Effects of chlamydia cell cultures on the proliferation of SARS-CoV in 293 cells

For the strain Chlamydia-1: four SARS-CoV single infection groups, two the strain Chlamydia-1single infection groups and one control group were cultured in the cell plate until the 6th day, then the cell cultures of two the strain Chlamydia-1 single infection groups and control group were collected. For two the strain Chlamydia -1 single infection groups, one groups’ culture mediums were collected, centrifuged at 12000 rpm for 1 min, and then its supernatants were collected, filtered with 0.22 um membrane, ready for use (extracellular cultures), while for the other group, both its culture mediums and cultured cells (by cell scraper) were collected, broken by ultrasonic wave on ice for ten times, 10 s/time, then centrifuged at 12000 rpm for 1 min, finally, the supernatants were collected, filtered with 0.22 um membrane, ready for use (extracellular + intracellular cultures). For the control group, its cell culture medium were collected, centrifuged at 12000 rpm for 1 min, and then the supernatant were collected, filtered with 0.22 um membrane, ready for use. Three SARS-CoV single infection groups were taken out and its growth mediums were discarded respectively, then, the above treated supernatants were added to them respectively. The above three groups were defined as extracellular group, extracellular + intracellular group and control group respectively, numbered as A_1_, B_1_, and C_1_. Finally, 200 uL 10% FBS DMKM were added to them and the remaining one SARS-CoV single infection group (numbered as D_1_) for continuing to incubate those groups until each detection. The viral load of each groups would be detected on the 6th, 7th, 8th and 9th day.

For the strain *C. psittaci*: the same as above, except replacing the strain Chlamydia-1 with the strain *C. psittaci*. The corresponding groups were number as A_2_, B_2_, C_2_ and D_2_, respectively.

### 1.4 Determination of IFUs

On the 3rd, 4th,5th, 6th 7th, 8th and 9th day, the round coverslips of each group were collected respectively, fixed in absolute methanol for 10 min, stained by Giemsa method for 45 min, then the number of IFUs would be calculated by inverted microscope under high power microscopic view. The number of IFUs was calculated by the following formula.

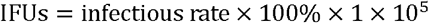

The meaning of the formula: the number of IFUs in cell cultures. Infectious rate is the number of IFUs divided by the number of cells. 1×10^5^ is the total number of cells.

### 1.5 Determination of viral load

Viral ribonucleic acid (RNA) was extracted by viral RNA kit (Qiagen, Germany), reversed by reverse transcription kit (Promega, United States). The sequence primers of SARS-CoV were synthesized by bio-engineering Co., Ltd. China (F:5’-GAAGCTATTCGTCACGTTCG-3’; R:5’-TAACCAGTAGGTACAGCTAC-3’). The viral complementary deoxyribonucleic acid (cDNA) was amplified and detected by SYBR Green PCR Master Mix kit (bao bio-engineering Co., Ltd, China) on the ABI7500 instrument (ABI, USA). For qRT-PCR reactions, the 20 uL reaction mixture included 2 uL cDNA, 10 uL 2×SYBR Green qPCR Mix, 0.4 uL of forward primer and 0.4 uL of reverse primer, and 7.2 uL RNAase-free water. Conditions for reactions were 55°C for 10 min and 95°C for 3 min, 45 cycles of amplification at 95°C for 10 s and 72 °C for 10 s.

### 1.6 Statistical Analysis

The data are presented as averages ± standard deviations (SDs), as indicated. Statistical comparisons were analyzed with two-tailed Student t-test and one-way analysis of variance (ANOVA) method. All of the statistical analyses were performed with statistical product and service solutions (SPSS) version 25.0. When p > 0.05, the results were not significant; when p < 0.05, the results were significantly different; when p < 0.01, the results were extremely significantly different.

## 2 Results

### 2.1 Results of co-infection of SARS-CoV and chlamydia in vitro

From figure 1A, we can see that the viral load of C_1_ group (from day 3 to day 9) and D_1_ group (form day 4 to day 9) has a extreme significant reduction compared to A_1_ group (P < 0.01). From figure 1B, we can see that the viral load of C_2_ group (from day 3 to day 9) and D_2_ group (from day 6 to day 9) has a significant reduction compared to A_2_ group (P < 0.05). From figure 1C, we can see that the viral load of C_1_ group (from day 3 to day 9), D_1_ group (from day 4 to day 9) and E_1_ group (from day 4 to day 9) has a extreme significant difference from that of C_2_ group, D_2_ group and E_2_ group respectively (P < 0.01).

**Figure 1.**
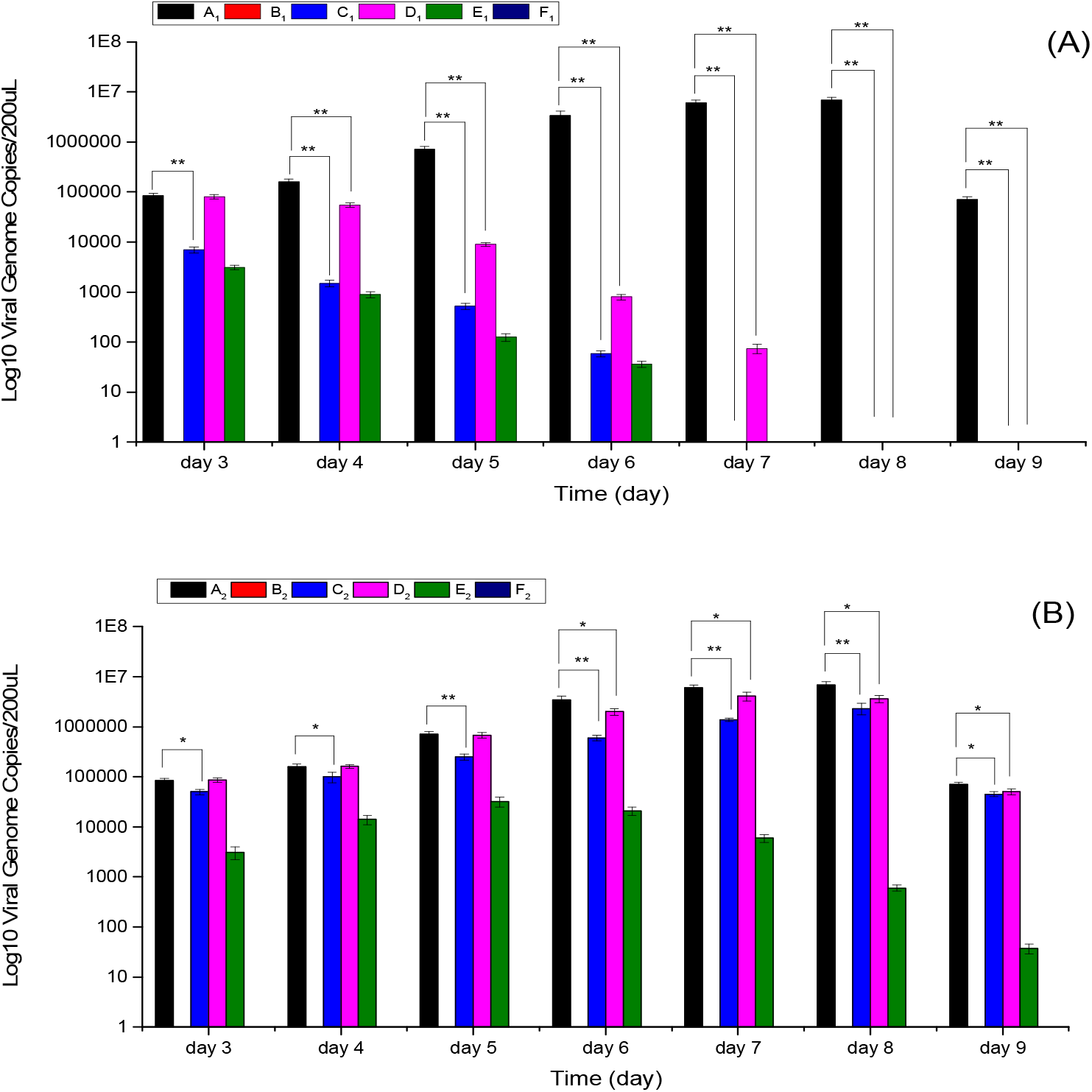
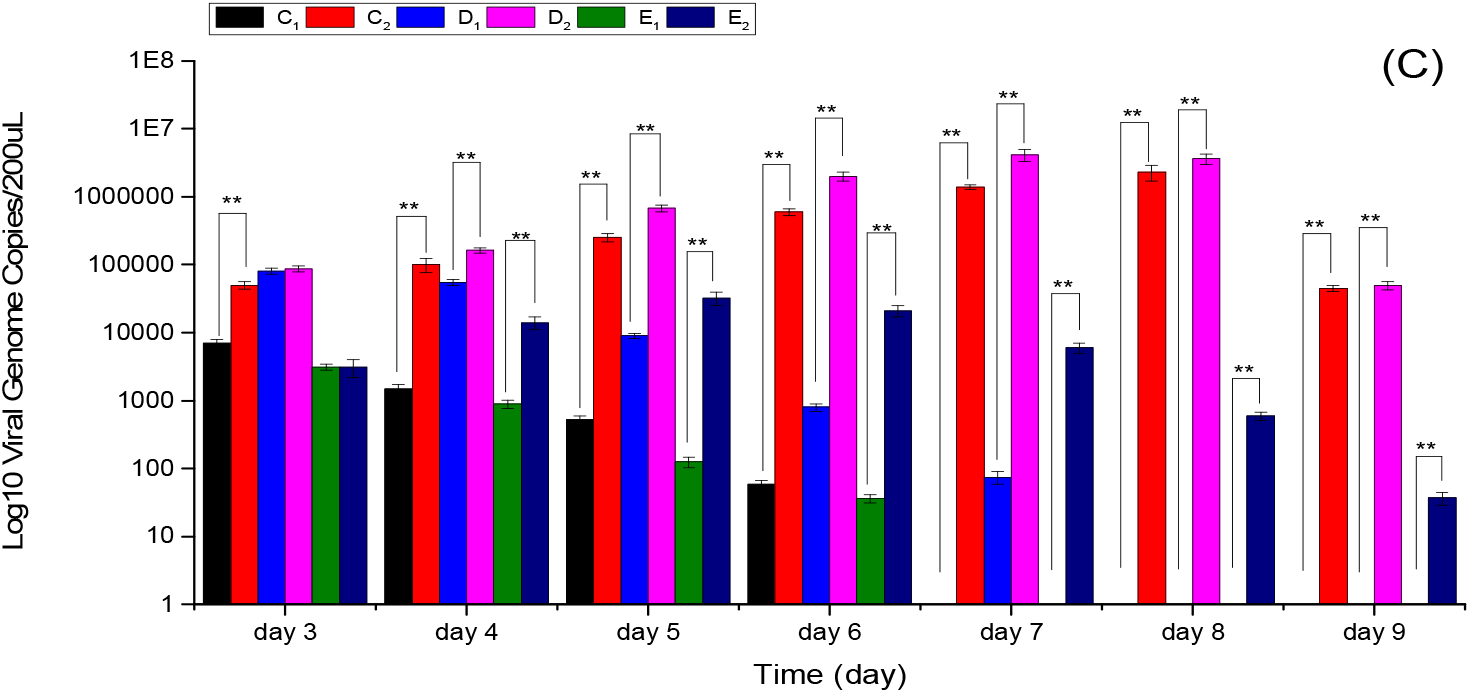
Time series changes of viral load in the co-infection of SARS-CoV and chlamydia in vitro. (A) Changes of viral load in the co-infection of SARS-CoV and the strain Chlamydia-1. (B) Changes of viral load in the co-infection of SARS-CoV and the strain *C. psittaci*. (C) Comparison of viral load between the co-infection of SARS-CoV and the strain Chlamydia-1 and the co-infection of SARS-CoV and the strain *C. psittaci*.

From figure 2A, we can see that the number of IFUs has no significant difference among B_1_ group, C_1_ group and E_1_ group (P > 0.05); It can be seen from figure 2B that the number of IFUs of C_2_ group (from day 3 to day 5 and day 7 to day 9) and E_2_ group (from day 4 to day 6) has a significant difference compared to B_2_ group (P < 0.05). Statistical significance between experimental groups was determined by the student’s t-test, as follows: *: p < 0.05, **: p < 0.01.

**Figure 2.**
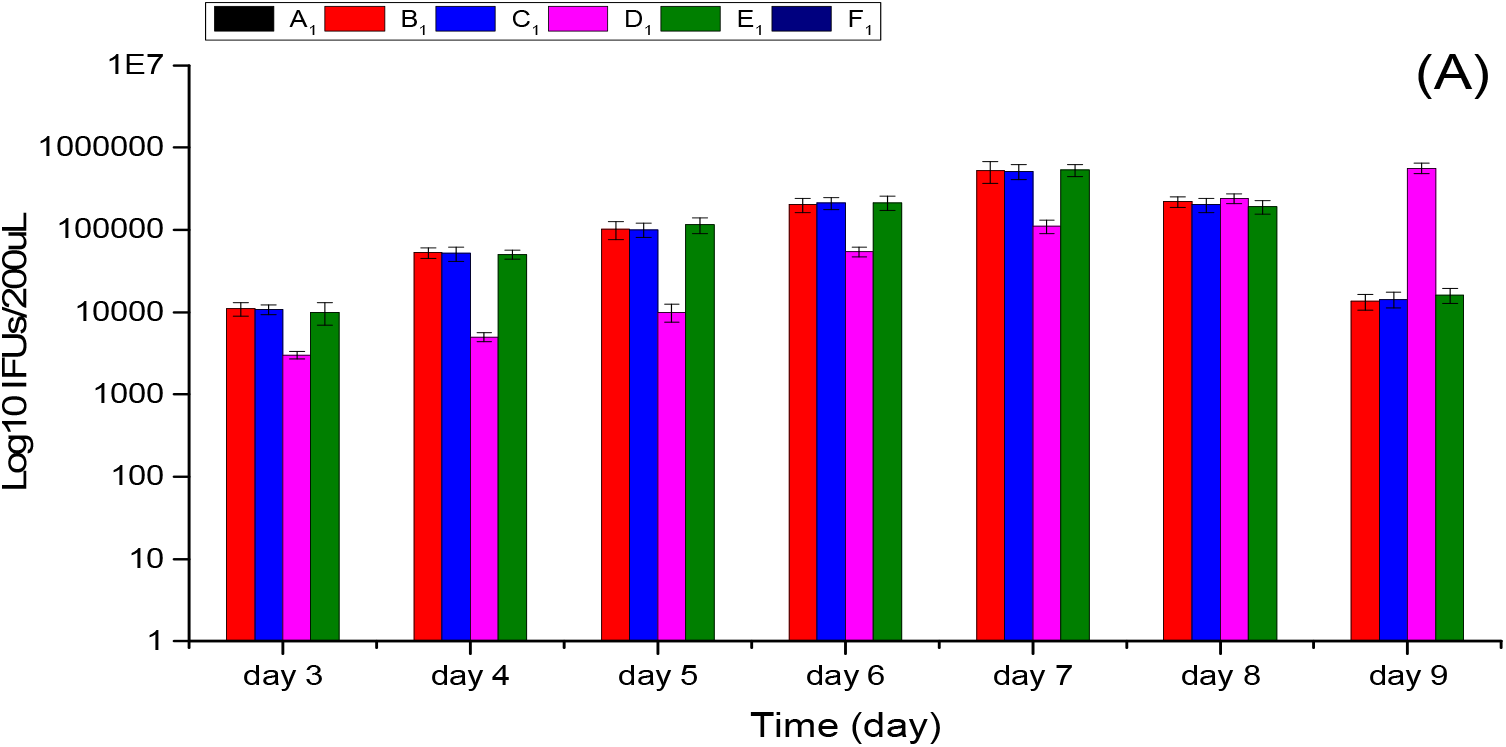
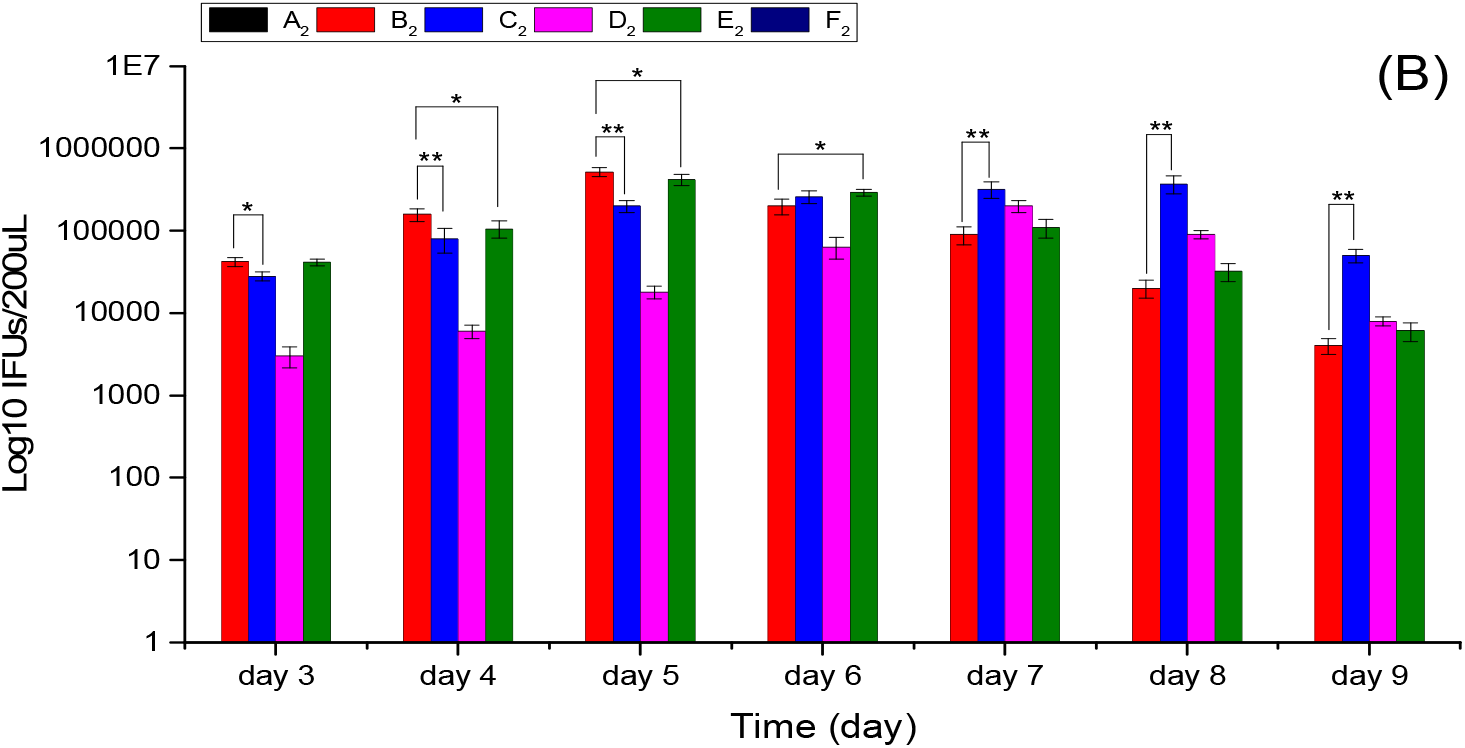
Time series changes of IFUs in the co-infection of SARS-CoV and chlamydia in vitro. (A) Changes of IFUs in the co-infection of SARS-CoV and the strain Chlamydia-1. (B) Changes of IFUs in the co-infection of SARS-CoV and the strain *C. psittaci*.

Therefore, the strain Chlamydia-1 has a extreme significant effect in decreasing the proliferation of SARS-CoV in vitro. while the strain *C. psittaci* seems also can significantly decrease the proliferation of SARS-CoV in vitro, its effect is significantly inferior to the strain Chlamydia-1. The proliferation of the strain Chlamydia-1 could not be affected by the proliferation of SARS-COV in vitro, while the proliferation of the strain *C. psittaci* seems to be affected by the proliferation of SARS-CoV in vitro.

### 2.2 Effects of chlamydia cell cultures on the proliferation of SARS-CoV in vitro

It can be seen from figure 3A that the viral load of A_1_ group (from day 7 to day 9) and B_1_ group (from day 7 to day 9) has a extreme significant reduction compared to C_1_ and D_1_ group (P < 0.01), and there is a extreme significant difference in reducing the viral load of SARS-CoV between A_1_ and B_1_ group (from day 7 to day 9) (P < 0.01). From figure 3B, we can see that the viral load of A_2_ group (from day 7 to day 8) and B_2_ group (from day 7 to day 8) has a significant reduction compared to C_2_ group and D_2_ group (P < 0.05). However, there is no significant difference in decreasing the viral load of SARS-CoV between A_2_ group and B_2_ group (from day 7 to day 8) (P > 0.05). From figure 3C, we can see that the viral load of A_1_ and B_1_ group has a extreme significant difference from that of A_2_ and B_2_ group respectively (from day 7 to day 9) (p < 0.01). Statistical significance among groups was determined by one-way ANOVA method.

**Figure 3.**
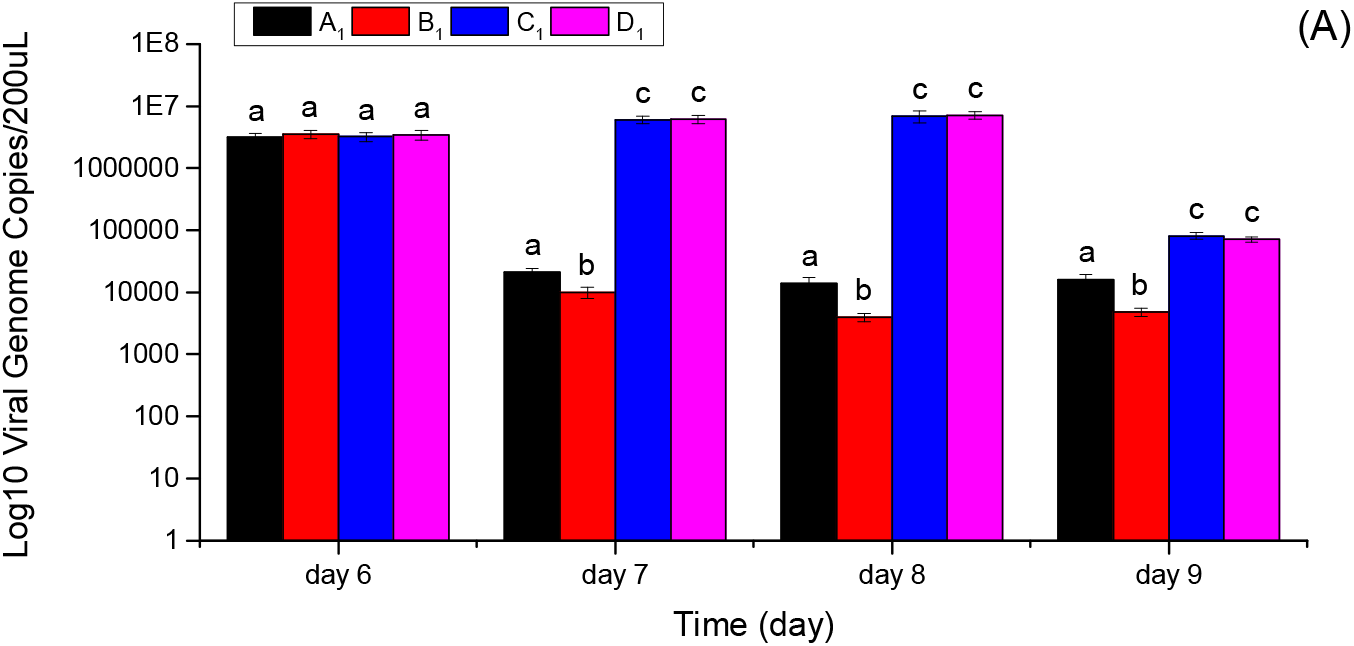
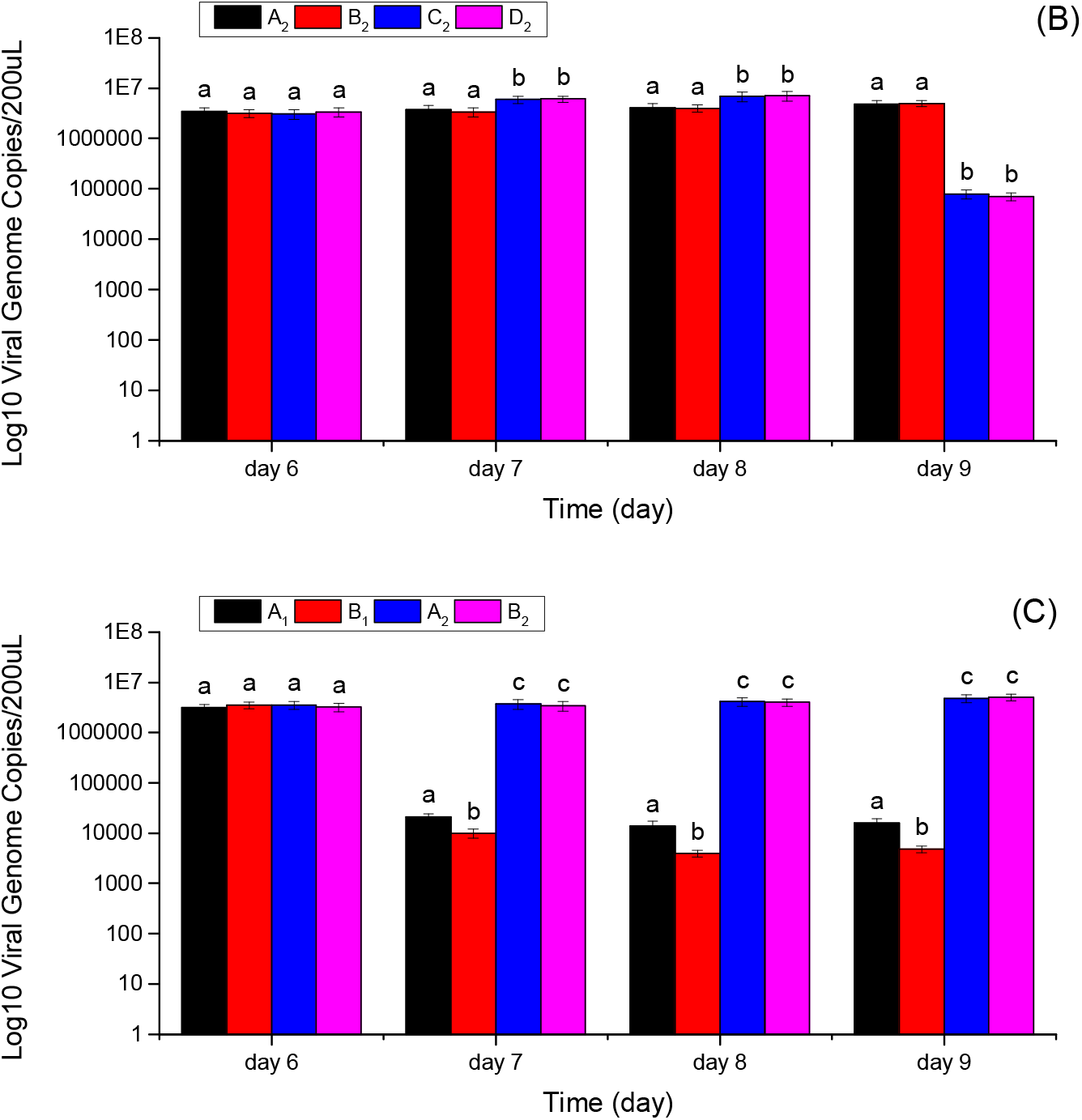
Effects of chlamydia cell cultures on the proliferation of SARS-CoV in vitro. (A) Effects of the strain Chlamydia-1 cell cultures on the viral load of SARS-CoV. (B) Effects of the strain *C. psittaci* cell cultures on the viral load of SARS-CoV. (C) Comparison of the effects between the Chlamydia-1 strain cell cultures and the strain *C. psittaci* cell cultures on the viral load of SARS-CoV. Bars with the same lower-case letters in two different columns show no significant difference (p > 0.05), and bars with different lower-case letters in two different columns show a significant difference (p < 0.05 or p < 0.01).

Therefore, foe the strain Chlamydia-1, it contains anti-SARS-CoV factors in its cell cultures, distributing both in the extracellular and intracellular cultures. While the strain *C. psittaci* cell cultures seems also has anti-SARS-CoV factor, its result is extremely significantly inferior to that of the strain Chlamydia-1 cell cultures. And for the strain *C. psittaci*, there is no significant difference between the extracellular + intracellular and extracellular cell cultures in decreasing the virus load of SARS-CoV.

## 3 Discussions

Because there was no SARS-CoV or few SARS-CoV was found in some tissue samples of SARS patients, but a large number of chlamydia particles were found in their tissue samples, in order to investigate the relationship between SARS-CoV and the related strain chlamydia in SARS patient’s autopsy issues, we conducted an experiment which involved the co-cultures of SARS-CoV and the related strain chlamydia^[17]^. We found that the strain chlamydia isolated from SARS patient’s autopsy issues could extremely significantly decrease the proliferation of SARS-CoV in vitro. It means that the strain chlamydia isolated from SARS-CoV patient’s autopsy issues may inhibit the proliferation of SARS-CoV, which benefits for exploring anti-SARS-CoV methods and drugs. Because there was no study that reported that some chlamydias could decrease the proliferation of related virus in vitro or vivo before that, and this work is only our preliminary observations, the specific mechanism that can explain the strain chlamydia isolated from SARS-CoV patient’s autopsy issues in decreasing the proliferation of SARS-CoV in vitro is still unknown. Therefore, in future works we will explore this aspects. Moreover, because our observed results indicated that the inhibitory factors distribute both in the extracellular and intracellular cultures, we will still try to identify and separate the inhibitory factors from the strain Chlamydia-1 cell cultures in future works. Finally, we noted that a recent study indicated that there was a lower likelihood of intensive care unit admission with bacterial co-infection in Coronavirus Disease-2019 (COVID-19) patients (r = -0.28, p=0.04), and *Chlamydia pneumoniae* was the most prevalent co-infecting bacteria found in 13 patients (28% in total 48 COVID-19 patients). Co-infection with *Chlamydia pneumoniae* has a negitive correlation with mortality in COVID-19 patients (r=-0.17).^[18]^ Combined with our study to analyze, we suggest that it should further explore the relationship between Severe acute respiratory syndrome coronavirus 2 and the strain *Chlamydia pneumoniae* in COVID-19 patients with the widely spread of COVID-19 across the world^[19]^.

## 4 Conclusions

We found that a strain chlamydia isolated from SARS patient’s autopsy issues could decrease the proliferation of SARS-CoV in vitro and the inhibitory factors distribute both in the extracellular and intracellular cultures, which benefits for us to further exploring the relationship between SARS-CoV and the related strain chlamydia in SARS patients, and even may provide helps for finding anti-SARS-CoV methods and drugs.

## Abbreviations

SARS-CoV: Severe acute respiratory syndrome coronavirus;
SARS: Severe acute respiratory syndrome;
IFUs: Inclusion-forming units;
HIV-1: Human immunodeficiency virus type 1;
*C. Psittaci*: *Chlamydia psittaci*;
TCID_50_: Fifty-percent tissue culture infectious dose;
FBS: Fetal bovine serum;
DMKM: Dulbecco’s modified Eagle’s medium;
PBS: Phosphate buffered saline;
qRT-PCR: Quantitative real-time polymerase chain reaction;
RNA: Ribonucleic acid;
cDNA: Complementary deoxyribonucleic acid;
SPSS: Statistical product and service solutions;
ANOVA: Analysis of variance;
COVID-19: Coronavirus disease-2019

## Acknowledgments

The authors are grateful to prof. Hong T for providing experimental virus, chlamydia, and cells and many helpful suggestions for improving the quality of this manuscript.

## Authors’ contributions

Conceived and designed the experiments: X.Q. Performed the experiments: X.Q., W.L. Analyzed the data: X.Q. W.L. Wrote the paper: X.Q. All authors read and approved the final manuscript.

## Availability of data and materials

The datasets used and/or analyzed during the current study are available from the corresponding author on reasonable request.

## Declarations

### Ethics approval and consent to participate

No applicable.

### Consent to publication

The authors declare they are consent for the manuscript publication.

